# Transcriptomic analysis reveals similarities in genetic activation of detoxification mechanisms resulting from imidacloprid and chlorothalonil exposure

**DOI:** 10.1101/327809

**Authors:** Justin Clements, Benjamin Sanchez-Sedillo, Christopher Bradfield, Russell L. Groves

## Abstract

The Colorado potato beetle, *Leptinotarsa decemlineata* (Say), is an agricultural pest of commercial potatoes in parts of North America, Europe, and Asia. Plant protection strategies within this geographic range employ a variety of pesticides to combat not only the insect, but also plant pathogens. Previous research has shown that field populations of *Leptinotarsa decemlineata* have a chronological history of resistance development to a suite of insecticides, including the Group 4A neonicotinoids. The aim of this study is to contextualize the transcriptomic response of *Leptinotarsa decemlineata* when exposed to the neonicotinoid insecticide imidacloprid, or the fungicides boscalid or chlorothalonil, in order to determine whether these compounds induce similar detoxification mechanisms. We found that chlorothalonil and imidacloprid induced similar patterns of transcript expression, including the up-regulation of a cytochrome p450 and a UDP-glucuronosyltransferase transcript, which are often associated with xenobiotic metabolism. Further, transcriptomic responses varied among individuals within the same treatment group, suggesting individual insects’ responses vary within a population and may cope with chemical stressors in a variety of manners. These results further our understanding of the mechanisms involved in insecticide resistance in *Leptinotarsa decemlineata*.

**Author Contribution Statement:** Conceived and designed the experiments: JC, CB, RLG. Performed the experiments: JC, BSS. Analyzed the data: JC, BSS. Wrote the paper: JC, BSS, RLG.

## Introduction

The development of insecticide resistance in the Colorado potato beetle (*Leptinotarsa deceminleata* (Say)) is a major concern for agricultural growers^1^. *L. decemlineata* are agricultural pests that originated in parts of North America and have now expanded their range to predominate in potato production areas of Europe and Asia^2^. Few natural enemies exist which can adequately control populations of *L. decemlineata* at manageable levels, and when unabated by plant protection strategies, populations can decrease agricultural potato yields by up to 100% through direct defoliation^2,3^. Populations of *L. decemlineata* have adapted from feeding on its native host plant, buffalobur (*Solanum rostratum*), to other agriculturally-relevant plants within the *Solanaceae* family, including potatoes, tomatoes, and eggplants^2^.

Insecticide classes such as organochlorines, organophosphates, synthetic pyrethroids, and carbamates have historically become ineffective for control of *L. decemlineata* ^4^ in many parts of their current geographic range and have been replaced by different chemical insecticides. Recent integrated pest management strategies for the control of *L. decemlineata* have often included the neonicotinoid insecticide, imidacloprid, a Group 4A nicotinic acetylcholine receptor agonist classified by the Insecticide Resistance Action Committee^5^. Imidacloprid is a water-soluble, systemically mobile insecticide often employed as an at-plant, in-furrow, or seed treatment application delivered at the beginning of the growing season. Imidacloprid selectively binds to an insect’s acetylcholine receptors on the post-synaptic membrane causing overstimulation, which leads to paralysis and death^6^. Since 2007, agriculturally-relevant populations of *L. decemlineata* have had measurable levels of resistance to imidacloprid^7,8^. Previous studies have shown that these populations originate from agricultural operations where insecticide resistance management (IRM) label guidelines of imidacloprid application have been followed. Unfortunately, even when IRM guidelines are adhered to, insecticide resistance may develop, and producers are forced to rotate their product choices to include different mode of action (MoA) classes for adequate control of this problematic pest species.

Comprehensive plant protection strategies must necessarily consider the control of other pest species, including fungal pathogens. In doing so, regular foliar applications of protectant fungicides are widely used and often occur throughout the growing season in potatoes when conditions or criteria for the risk for disease development have been surpassed. Further, fungicide applications may be applied in a tank mixture with, or may follow, neonicotinoid insecticide (e.g. imidacloprid) application leading to combined or even subsequent exposure to both the fungicide and insecticide. One of the most widely used broad spectrum, and multi-site MoA fungicides used in potatoes is chlorothalonil. Chlorothalonil is classified by the Fungicide Resistance Action Committee as a multi-site MoA, chloronitrile (phthalonitrile)^9,10^, and may be applied weekly during periods of high disease risk as a wide range of formulated products applied at rates of 0.67 kg/ha of active ingredient per acre^11^. Another commonly used, single-site MoA fungicide applied to potatoes is the carboxamide (MoA Group 7) boscalid. Here again, potatoes can receive 2-4 successive, foliar applications of boscalid as a protectant fungicide at rates of 0.49 kg/ha^12^. In the context of an organism’s continued exposure to both insecticides and fungicides over a crop season, populations of *L. decemlineata* may become tolerant to a normally toxic chemical when previously exposed to a different chemical MoA that is similarly metabolized by identical proteins; operationally defined here as cross-resistance. Drivers of insecticide cross-resistance have been characterized in many pest species including, *Cydia pomonella*^13^, *Aedes aegypti*^14^, and *Bactrocera dorsalis*^15^. Clements et al. (2017) previously demonstrated that imidacloprid tolerance can be partially ablated through the RNA interference of transcripts encoding for a cytochrome p450, suggesting the major role that these proteins play in resistance within select populations^16^. Recent transcriptomic investigations have revealed differentially expressed transcripts between imidacloprid resistant and susceptible populations of *L. decemlineata*, and suggest that cytochrome p450s, cuticular proteins, ABC transporters, and glutathione related proteins are up-regulated in resistant populations^8^. Still other investigations in multiple insect taxa further support these findings and suggest that a discrete set of genes appear to be responding to insecticides ^16,17,18^, suggesting that these proteins may be playing vital roles in xenobiotic metabolism. We hypothesize that fungicides may be able to up-regulate similar genetic detoxification mechanisms as imidacloprid, and therefore, could play a role in conferring cross-resistance.

The current study employed RNA sequencing to characterize the transcriptomic response of *L. decemlineata* when exposed to imidacloprid, chlorothalonil, or boscalid in controlled investigations, and investigated whether similar patterns of genetic expression of detoxification mechanisms were induced by these compounds. We compared the transcriptomic profiles of known genes involved in the detoxification of imidacloprid to those that were up-regulated in response to field-relevant application rates of both boscalid and chlorothalonil. The up-regulation of these genes when exposed to boscalid or chlorothalonil may confer cross-resistance to neonicotinoid insecticides.

## Materials and Methods

### Data Availability

All relevant data are contained within the paper and its supporting information files. The accession numbers for the raw RNA sequencing data are deposited in the Sequence Read Archive (SRA) SRP144770 and The Transcriptome Shotgun Assembly project has been deposited at DDBJ/EMBL/GenBank under the accession GGNV00000000. The version described in this paper is the first version, GGNV01000000.

### Ethics Statement

No specific permits were required for field collection or experimental treatment of *L. decemlineata* for the study described.

### Insect Maintenance

Approximately 300 adult beetles were initially collected on June 20^th^, 2016 from the Arlington Agricultural Research Station, Arlington, Wisconsin (AARS, 43.315527, −89.334545), where populations have little prior exposure to insecticides, and this population remains highly susceptible to imidacloprid^7,8^. Healthy adult beetles were hand-collected from the canopy of potato plants, placed in plastic containers, and returned to the University of Wisconsin-Madison. Reproducing populations of *L. decemlineata* were sustained on healthy potato plants in mesh cages under a 16:8 hour light:dark (L:D) photoperiod. Untreated foliage from potato plants was obtained from plants grown at the University of Wisconsin-Madison greenhouse and provided to beetles daily. Adult beetles were given the opportunity to randomly mate and lay clutches of eggs on potato foliage. Egg masses were collected daily and placed on filter paper in 100 × 15 mm petri dishes (Corning, Corning, New York) and held at 26°C, 70% relative humidity (RH), and 16:8 (L:D) photoperiod. Following egg hatch, larvae were provided untreated foliage daily and maintained as cohort groups throughout the remainder of their larval development before being returned to mesh cages with fresh potato plants and soil and allowed to pupate and subsequently emerge as adults.

### Insecticide and Fungicide Feeding Exposure

From the previously described laboratory colony, twenty-four, 7^th^ generation adult beetles (emerging from within 24 hours of pupation) were removed, and these selected adult beetles were used in the following experiments. Adult beetles were placed in individual 100 × 15 mm Petri dishes (Corning, Corning, NY,) on top of filter paper, and starved for 24 hours in a Percival incubator (Percival Scientific, Perry, IA) at 26°C, 70% humidity, and 16:8 (L:D) in advance of any experiments. Fresh potato leaves were removed from healthy, greenhouse-grown plants and cleaned of any debris prior to the excision of a 2.01cm^2^ leaf disk. For each chemical compound assayed, one leaf disk for each individual beetle was dipped in acetone solutions containing field-relevant concentrations of technical chlorothalonil (6.9 μg/μl), boscalid (13 μg/μl), imidacloprid (0.000079 μg/μl), or an acetone control (n=6 insects/treatment). Adult beetles were presented individual leaf disks in their petri dish. Beetles were monitored until the insect had located and started feeding upon their leaf disks. Once feeding was initiated on the leaf disk, the adults were then given 24 hours to consume the entire leaf disk and were held at 26°C, 70% humidity, and 16:8 (L:D) throughout this interval. After 24 hours, we randomly selected four individual adults representing each compound from among those which had consumed the entire leaf disk provided, and these individuals were later used for transcriptomic analyses.

### RNA Extraction, RNA Sequencing and Transcriptome Assembly

Sixteen adult *L. decemlineata*, 4 from each treatment (boscalid, chlorothalonil, imidacloprid, and control), were sacrificed exactly 24 hours after the onset of leaf ingestion. Total RNA was extracted from each adult beetle using Trizol (Life Technology, Grand Island, NY). DNA contamination was removed with TurboDNase (Life Technology, Grand Island, NY) and total RNA was purified through an EtOH precipitation, air dried until no visible liquid was observed, and then suspended in 50 μL DNase/RNase-free H_2_O. Quality of RNA was initially assessed using a Nanodrop (ThermoFisher Scientific, Waltham, MA). From each sample, 2500 ng of total RNA was provided to the University of Wisconsin-Madison Biotechnology Center (UWMBC). RNA quality was further examined using a 2100 Bioanalyzer (Agilent Technologies, Santa Clara, CA). The UWMBC was also contracted to isolate and generate mRNA libraries and subsequently perform Illumina HiSeq 2×125bp sequencing (Illumina, San Diego, CA). From raw RNA sequencing reads, the UWMBC generated de novo transcriptome assembly using Trinity bioinformatics software. A Busco (Version 3.0.2) analysis was conducted on the assembled transcriptome to characterize coverage.

### Differentially-Expressed Transcripts and Transcriptome Analysis

Transcript expression was determined using Trinity bioinformatics software (RSEM). TPM (transcripts per million) were calculated for each contig in the transcriptome. Further, differentially-expressed transcripts were determined using edgeR, which produced a log fold change along with the corresponding FDR (false discover rate). In the analysis, transcripts were considered to be over-or under-expressed with a log_2_ fold change greater than 2 and an FDR value < 0.05. A sample correlation matrix was generated for differentially-expressed transcripts (**Supplemental Figure S1**). Three differentially-expressed transcript analyses were conducted, including transcript abundance between the control group and each of the chemical induction groups. Further, from the sample correlation matrix, it was noted that the transcript expression of specific beetle replicates (Imidacloprid_rep2, Imidacloprid_rep3, and Chlorothalonil_rep1) were statistically similar, but distinct from the remaining experimental individuals. These statistically similar individuals were classified as the “Effect” group, while the remaining individuals were classified as the “No Effect” group. A differential expression analysis was conducted between Imidacloprid_rep2, Imidacloprid_rep3, and Chlorothalonil_rep1 (Effect) and all other experimental individuals (No Effect). Trinity transcripts were classified using BLAST standalone. Reference protein sequences (Refseq) from all Coleoptera were downloaded from NCBI for a total of 119,760 sequences. A reference database was created with the protein sequences for BLAST analyses and comparison. Using BLAST standalone with BLASTx, the total unique trinity contigs in the transcriptome were compared to the reference proteins (*E* value <10^−3^). Transcripts were classified based on the NCBI nomenclature returned by BLASTx. BLASTx results were uploaded into Blast2Go^19^ for further data analysis. BLASTx was conducted on the entire transcriptome. Within the analysis, we chose to focus on the longest isoform of each Trinity gene set. Components were mapped to the corresponding GO terms, then the annotation step was run with a cutoff of E_value_ < 1E-3, annotation cut off > 45, and GO weight >5.

### Confirmation of Transcript Abundance with Quantitative PCR

To confirm the patterns of transcript abundance, four differentially-expressed targets were chosen from the transcriptome. From the purified RNA, cDNA was generated using a Superscript III kit (ThermoFisher Scientific, Waltham, MA). The cDNA was diluted to a final concentration of 5ng/μl of RNA equivalent input for qPCR. The ribosomal protein 4 (RP4)^20^ was used as a reference gene for all analyses. The qPCR reaction was run on a CFX-96 platform (Bio-Rad Laboratories, Hercules, CA) with a master mix of Bullseye EverGreen (MIDSCI, Valley Park, MO). Primer and primer efficiency can be found in **Supplemental Table S1**. Primer specificity was checked against the transcriptome using BLAST. Primer efficiencies were calculated and optimized. Triplicate reactions were run at 95°C for 10 min followed by 95°C for 15 s, 62°C for 60 s, for 40 cycles. Data were collected for each biological replicate, and relative expression of resistant strains to susceptible strains was calculated using the Pfaffl methodology^21^.

### Complimentary Chlorothalonil and Imidacloprid Assays Conducted to Verify Transcripts of Interest

The insecticide and feeding exposure assay to chlorothalonil and imidacloprid was repeated twice more to further validate the genetic response described by the transcriptomic analysis. RNA was extracted and cDNA was generated to observe genetic induction of similar detoxification mechanisms (n=4 insects/compound/trial 2, and n=5 insects/compound/trial 3). Quantitative PCR was conducted as previously stated to observe the expression for each exposure group to the transcripts of interest (cytochrome P450 (6k1 isoform X1) and UDP-glucuronsyltransferase (2b7-like)).

## Results

### Transcriptome Assembly

Illumina short-read sequences from mRNA isolated from 16 adult *L. decemlineata*, 4 from each treatment (boscalid, chlorothalonil, imidacloprid, and control), were compiled into a transcriptome using Trinity bioinformatics software^22^. The compiled transcriptome revealed 418,015 total transcripts and 218,882 unique trinity ‘genes’. When compared to the compiled NCBI data base of reference Coleoptera sequences, there were 55,064 total transcripts with a BLASTx match. Further, a Busco analysis (Version 3.0.2) conducted on the transcriptome revealed that 97 percent coverage was achieved. Trinity Transcriptome Assembly Statistics can be found in Table 1.

**Table 1:** Trinity De Novo Assembly Statistics.

| Trinity Transcriptome Assembly Statistics |  |
| --- | --- |
| Total Trinity Transcripts | 418,015 |
| Total Trinity 'Genes' | 218,882 |
| BLASTx Hits (e value ≥ 0.001) | 55,064 |
| Percent GC | 36.53 |
| Contig N50 | 1,074 |
| Median Contig Length | 349 |
| Average Contig | 661.56 |
| Total Assembled Bases | 276,543,627 |
| Busco Analysis of Transcriptome |  |
| C:97.0%[S:38.1%,D:58.9%],F:2.5%,M:0.5%,N:1658 |  |
| 1609 Complete BUSCOs (C) |  |
| 632 Complete and single-copy BUSCOs (S) |  |
| 977 Complete and duplicated BUSCOs (D) |  |
| 42 Fragmented BUSCOs (F) |  |
| 7 Missing BUSCOs (M) |  |
| 1658 Total BUSCO groups searched |  |

### Differential Expression Analysis

Several classes of previously described enzymatic detoxification mechanisms were up or down-regulated in response to both the insecticide or fungicide treatments, as listed in Table 2. Differentially-expressed transcripts were classified using a log_2_ fold change ≥ 2 and an FDR < 0.05 as seen in Table 2. The specific classes of detoxification mechanisms examined in the current study were based on previously classified detoxification mechanisms^1,8,18,23^.

**Table 2:**
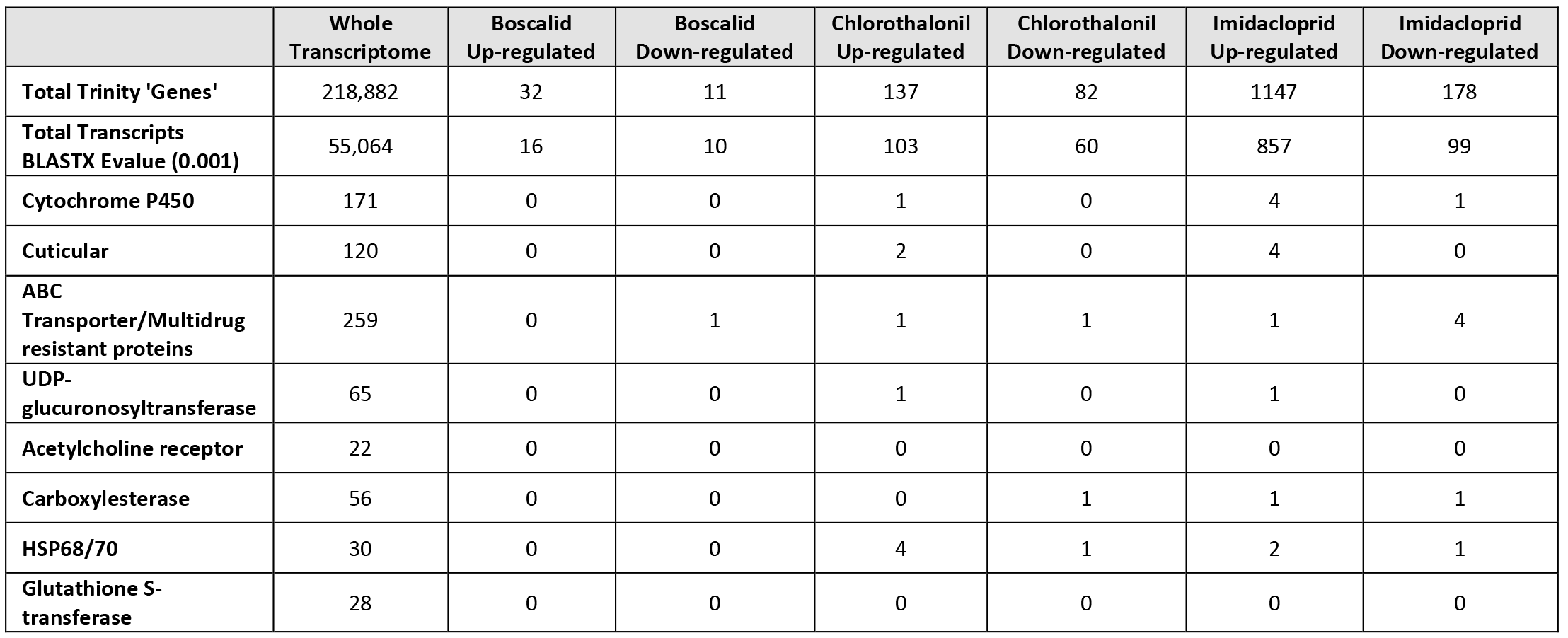
Differentially expressed transcripts between the control adult beetles and fungicide and insecticide exposures. Differentially-expressed transcripts were classified using a log2 fold change > 2 and an FDR < 0.05

|  | Whole Transcriptome | Boscalid Up-regulated | Boscalid Down-regulated | Chlorothalonil Up-regulated | Chlorothalonil Down-regulated | Imidacloprid Up-regulated | Imidacloprid Down-regulated |
| --- | --- | --- | --- | --- | --- | --- | --- |
| Total Trinity 'Genes' | 218,882 | 32 | 11 | 137 | 82 | 1147 | 178 |
| Total Transcripts BLASTX Evalue (0.001) | 55,064 | 16 | 10 | 103 | 60 | 857 | 99 |
| Cytochrome P450 | 171 | 0 | 0 | 1 | 0 | 4 | 1 |
| Cuticular | 120 | 0 | 0 | 2 | 0 | 4 | 0 |
| ABC Transporter/Multidrug resistant proteins | 259 | 0 | 1 | 1 | 1 | 1 | 4 |
| UDP-glucuronosyltransferase | 65 | 0 | 0 | 1 | 0 | 1 | 0 |
| Acetylcholine receptor | 22 | 0 | 0 | 0 | 0 | 0 | 0 |
| Carboxylesterase | 56 | 0 | 0 | 0 | 1 | 1 | 1 |
| HSP68/70 | 30 | 0 | 0 | 4 | 1 | 2 | 1 |
| Glutathione S-transferase | 28 | 0 | 0 | 0 | 0 | 0 | 0 |

Examination of differentially-expressed transcripts revealed several previously described insecticide detoxification mechanisms, including multiple cytochrome p450s and cuticular proteins. Further examination of the similarities between imidacloprid or chlorothalonil exposure revealed two known enzymatic mechanisms of insecticide detoxification to be statistically overexpressed in both the imidacloprid and chlorothalonil treatment groups. These previously described pesticide detoxification mechanisms included a cytochrome P450 **(6k1 isoform X1)** and UDP-glucuronsyltransferase **(2b7-like**) that were independently overexpressed as a result of both treatments. Classified enzymatic detoxification mechanisms based on log_2_ fold change and FDR can be found in Table 3. Examination of differentially-expressed transcripts between boscalid exposure groups and the control group revealed no statistically significant change in transcript expression for known detoxification mechanisms.

**Table 3:** Up-regulated transcripts that could encode for known pesticide detoxification mechanisms induced by either imidacloprid or chlorothalonil.

| NCBI Blast Match | Trinity Transcript ID | Log<sub>2</sub>FC | FDR |
| --- | --- | --- | --- |
| ***Imidacloprid Induction group*** |  |  |  |
| Probable cytochrome P450 | DN45930_c0_g1 | 10.24 | 8.47E-05 |
| Heat shock 70 kDa cognate 4-like | DN45929_c0_g1 | 9.30 | 0.00027 |
| Endocuticle structural glyco bd-8-like | DN53725_c1_g1 | 6.29 | 0.0025 |
| **Cytochrome P450 6k1 isoform X1** | **DN61141_c1_g1** | **5.86** | **0.0032** |
| Pupal cuticle 20-like | DN59030_c2_g1 | 5.84 | 0.010 |
| Endocuticle structural glyco bd-5-like | DN23859_c0_g1 | 5.70 | 0.0099 |
| Cuticle-like | DN48928_c1_g1 | 4.79 | 0.011 |
| Cytochrome P450 4d2-like | DN46083_c0_g3 | 4.69 | 0.0019 |
| **UDP-glucuronosyltransferase 2B7-like** | **DN61595_c0_g3** | **3.99** | **0.017** |
| Probable cytochrome P450 49a1 | DN47979_c8_g1 | 3.96 | 0.0024 |
| heat shock 68-like | DN54580_c0_g1 | 3.13 | 0.017 |
| Probable multidrug resistance-associated lethal(2)03659 | DN63738_c2_g1 | 2.82 | 0.0039 |
| Venom carboxylesterase-6-like | DN48501_c1_g1 | 2.82 | 0.038 |
| ***Chlorothalonil Induction Group*** |  |  |  |
| Uncharacterized oxidoreductase | DN44684_c0_g1 | 5.08 | 0.046 |
| **Cytochrome P450 6k1 isoform X1** | **DN61141_c1_g1** | **4.45** | **0.014** |
| Heat shock 70 A1-like isoform X1 | DN62524_c2_g4 | 4.43 | 0.00013 |
| Heat shock 68a | DN62524_c2_g1 | 4.16 | 1.46E-07 |
| Heat shock 70 A1-like isoform X1 | DN62524_c2_g2 | 3.91 | 1.21E-06 |
| **UDP-glucuronosyltransferase 2B7-like** | **DN61595_c0_g3** | **3.89** | **0.046** |
| Heat shock 70 A1 | DN41892_c0_g1 | 3.55 | 0.00082 |
| Pupal cuticle 20-like | DN44960_c0_g1 | 3.45 | 0.037 |
| Cuticle 7-like | DN45742_c0_g1 | 2.96 | 0.047 |
| Probable multidrug resistance-associated lethal(2)03659 | DN63738_c2_g1 | 2.59 | 0.023 |

Highlighted transcripts are significantly up-regulated in both imidacloprid and chlorothalonil groups

When a correlation matrix was generated between all individuals within the experimental and control groups, a unique distribution of overall transcript expression among test organisms was observed (**Supplemental Figure S1)**. Transcript expression patterns for the specific adult beetles: Imidacloprid_2, Imidacloprid_3 and Chlorothalonil_1, significantly grouped together (Effect) from the rest of the test individuals (No Effect). Differentially-expressed transcripts were examined between these two groups and patterns are illustrated in Tables 4 and 5.

**Table 4:**
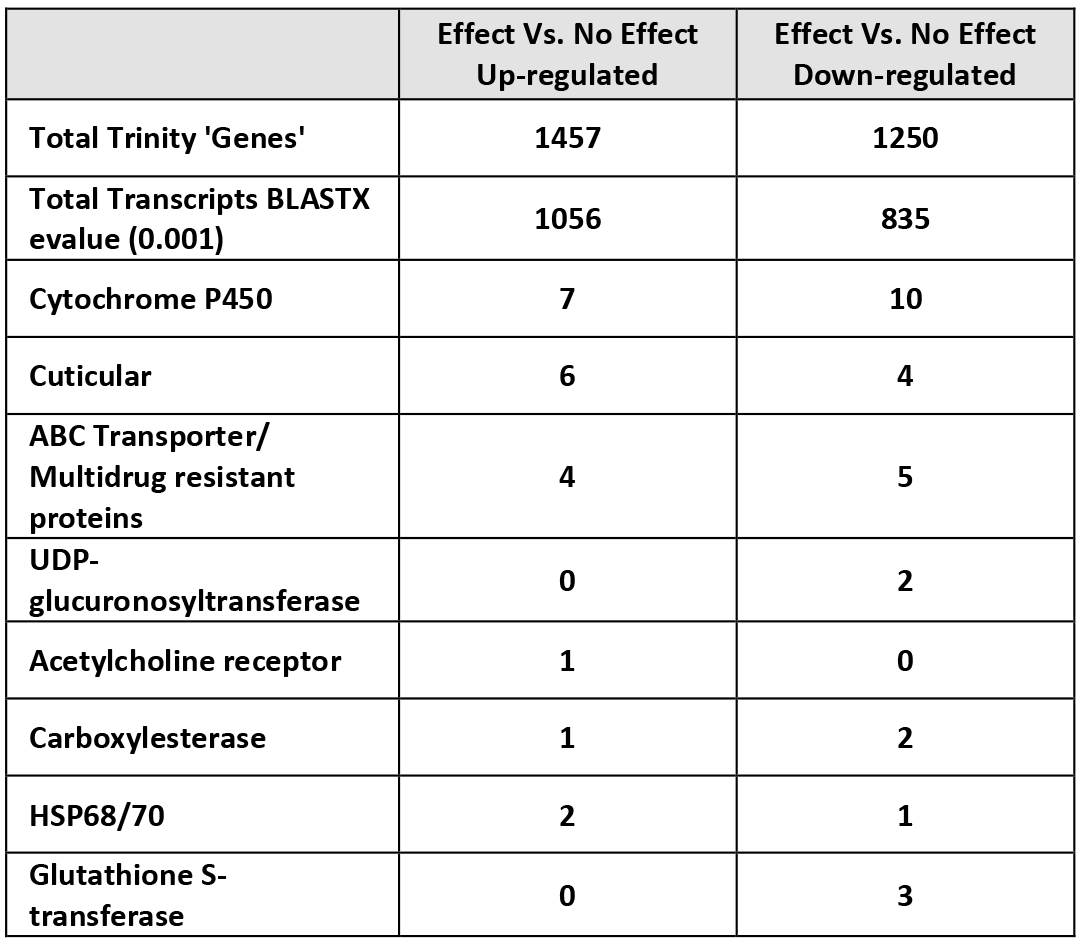
Differentially-expressed transcripts between Effect and No Effect groups.

**Table 5:** Up-regulated transcripts that could encode for known pesticide detoxification mechanisms induced in the Effect group as compared to the No Effect group.

| NCBI Blast Match | Trinity Transcript ID | Log<sub>2</sub>FC | FDR |
| --- | --- | --- | --- |
| **Effect Vs. No Effect** |  |  |  |
| Acetylcholine receptor subunit alpha-like | DN52191_c2_g3 | 11.70 | 2.96E-96 |
| Heat shock 70 kDa cognate 4-like | DN45929_c0_g1 | 10.68 | 1.99E-118 |
| Probable cytochrome P450 | DN45930_c0_g1 | 10.55 | 6.37E-157 |
| NADH-quinone oxidoreductase subunit B 2 | DN33393_c0_g1 | 9.34 | 9.63E-46 |
| Endocuticle structural glyco bd-8-like | DN53725_c1_g1 | 6.88 | 2.30E-18 |
| Cuticle 7-like | DN42933_c0_g1 | 6.62 | 6.28E-14 |
| Endocuticle structural glyco bd-5-like | DN23859_c0_g1 | 6.38 | 3.40E-22 |
| Pupal cuticle 20-like | DN59030_c2_g1 | 6.28 | 1.36E-21 |
| Cytochrome P450 4d2-like | DN46083_c0_g3 | 6.28 | 7.77E-56 |
| Cuticle-like | DN48928_c1_g1 | 6.12 | 2.58E-28 |
| Multidrug resistance-associated 1 isoform X3 | DN52951_c2_g1 | 5.63 | 4.28E-11 |
| Probable cytochrome P450 4d14 | DN46083_c0_g2 | 5.43 | 1.55E-11 |
| Cytochrome P450 6k1-like | DN43906_c0_g1 | 4.50 | 2.03E-06 |
| Heat shock 68-like | DN54580_c0_g1 | 4.12 | 7.52E-41 |
| Probable cytochrome P450 49a1 | DN47979_c8_g1 | 3.90 | 2.71E-12 |
| Probable multidrug resistance-associated lethal(2)03659 isoform X1 | DN48864_c1_g1 | 3.02 | 0.00013 |
| Carboxylesterase 5A | DN56141_c0_g1 | 2.73 | 0.0026 |
| Probable cytochrome P450 305a1 | DN51839_c1_g1 | 2.47 | 5.83E-05 |
| Probable multidrug resistance-associated lethal(2)03659 | DN63738_c2_g1 | 2.43 | 3.10E-06 |
| Pupal cuticle 20-like | DN44960_c0_g1 | 2.40 | 0.0048 |
| Cytochrome P450 4C1-like isoform X1 | DN45995_c0_g1 | 2.33 | 2.61E-05 |
| Multidrug resistance-associated 4-like | DN48293_c3_g1 | 2.06 | 9.42E-19 |

Examination of over-expressed transcripts within the Effect group revealed previously described detoxification mechanisms, including 7 cytochrome p450’s, multiple cuticular proteins, multidrug resistant proteins, and a highly up-regulated acetylcholine receptor subunit (alpha-like). Enrichment analysis on all the significantly up-regulated transcripts was conducted for each treatment group including the Effect vs No Effect group and can be found as **Supplemental Table S2**.

### Quantitative PCR Confirmation

Quantitative PCR confirmed transcript expression obtained from the RNA-sequencing data. A subset of differentially-expressed transcripts was confirmed, validating the RNA-sequencing and transcriptomic data. Further, separately from the transcriptomic study, two additional groups (Trial 2 and 3) of adult beetles were exposed to imidacloprid or chlorothalonil to examine if a similar genetic response for the two classified enzymatic detoxification mechanisms (cytochrome P450 6k1 isoform X1 and UDP-glucuronsyltransferase 2b7-like) was induced (Table 6). Threshold cycle values along with their standard deviations can be found in **Supplemental Table S3.**

**Table 6:** Differentially-expressed transcript values as reported from edgeR and quantitative PCR. Values represent a log_2_ transformation.

|  | Cytochrome P450<br>6k1 isoform X1<br>(DN61141) | UDP-glucuronosyltransferase<br>2B7-like<br>(DN61595) | Acetylcholine receptor<br>subunit alpha-like<br>(DN52191) | Probable<br>cytochrome P450<br>(DN45930) |
| --- | --- | --- | --- | --- |
| Transcriptome Chlorothalonil (TMM) | 5.86 | 4.45 | NA | NA |
| Validation of Chlorothalonil RNA in<br>Transcriptome (qPCR) | 4.97 | 2.91 | NA | NA |
| Transcriptome Imidacloprid (TMM) | 3.99 | 3.85 | NA | NA |
| Validation of Imidacloprid RNA in<br>Transcriptome (qPCR) | 2.88 | 3.22 | NA | NA |
| Transcriptome Effect (TMM) | NA | NA | 11.7 | 10.5 |
| Validation of Effect group RNA in<br>Transcriptome (qPCR) | NA | NA | 12.4 | 10.11 |
| Trail 2 rerun exposure to<br>chlorothalonil | 5.23 | 3.52 | NA | NA |
| Trail 2 rerun exposure to<br>imidacloprid | 1.17 | 0.45 | NA | NA |
| Trail 3 rerun exposure to<br>chlorothalonil | 3.07 | 6.47 | NA | NA |
| Trail 3 rerun exposure to<br>imidacloprid | -1.35 | 1.82 | NA | NA |

## Discussion

The ability of the Colorado potato beetle to develop insecticide resistance to multiple classes of pesticides has been well documented^1,4,8^ What is not immediately apparent from these investigations are the genetic mechanisms by which these insects adapt, survive and establish resistance. Further, the ability of one chemical (precursor) to activate a resistance mechanism that, in turn, can detoxify other relevant pesticides has also been established and has operationally been described as cross-resistance^24^. In the current manuscript, we hypothesize that a select set of agri-chemicals commonly applied in the potato agro-ecosystem and targeting unrelated pest taxa (fungi versus insects), can activate similar genetic mechanisms of detoxification and potentially have inadvertent consequences on *L. decemlinaeata* susceptibility to insecticides. Here we establish the genetic response in the form of transcript induction of known enzymatic detoxification mechanisms under lab conditions to the pesticides imidacloprid, chlorothalonil, and boscalid. Imidacloprid has been established as a highly effective insecticide used to control populations of *L. decemlineata*, while chlorothalonil and boscalid are fungicides used to control foliar pathogens of potato including *Phytophthora infestans* or *Alternaria solani,* (late and early blight of potato, respectively). Undoubtedly, populations of *L. decemlineata* in many potato producing areas of the United States are regularly exposed to repeat applications of all three compounds, especially those populations in the Midwest and Eastern production areas where the threat of foliar disease is greatest^1,9^. Our focus here, however, is not to demonstrate cross-resistance between fungicides and insecticides, but rather to demonstrate that selected insecticides and fungicides can induce a similar genetic detoxification response, which may increase the potential for cross-resistance in a field setting.

A transcriptome generated from total RNA representing 16 adult *L. decemlineata* was created. These individuals represented experimental groups treated with different pesticides, including imidacloprid, chlorothalonil, or boscalid. These compounds were selected based upon their frequency of use and regular occurrence in potato. Transcripts were compared to reference Coleoptera protein sequences from NCBI to determine their identity, revealing a total of 218,882 unique transcripts and 55,065 transcripts that had a corresponding BLASTx match. Using the transcriptome, its corresponding transcripts, and the RNA sequencing data, we determined differentially-expressed transcripts. Differentially-expressed transcripts were generated for each pesticide exposure group verses the control. In our analysis, we focused on the classification of previously known mechanisms of pesticide resistance. Toxicokinetic and toxicodynamic mechanisms are two broad categories of protection that organisms possess to respond to toxins^25^. Toxicokinetic mechanisms describe the biotransformation and excretion of the toxicant (insecticide) after it has been absorbed into the body. A toxicodynamic driver of resistance includes selection of target site insensitivity, such as a change in the peptide residues at the site of action, which lowers the affinity for the target, thus lowering the effective concentration to produce an adverse effect^26^. Toxicodynamic response describes the interaction of the toxicant at the target site of action, and the molecular alterations within the cell that elicit an adverse effect. Toxicokinetic mechanisms include the enhancement of metabolic processes that break down the chemical into subsequent metabolites and transport the xenobiotic away from sites of sensitivity. The phase 1 and phase 2 metabolism of neonicotinoids has been well characterized and has identified enzymatic proteins that have been linked to the detoxification of imidacloprid, which include cytochrome P450s, monooxygenases, and glutathione related proteins^18,23^. The transport of a toxic chemical away from the target site of action is partly achieved through the use of ABC transporters^17^. Additionally, transport can be facilitated for the sequestration of the chemical into non-metabolically active tissues such as the cuticle to make them inert ^24,27^. A third mechanism involved in resistance is behavioral modification in insects’ phenology, habitat choices, or foraging behavior^1^.

From the toxicokinetic and toxicodynamic mechanisms, we chose to focus on phase 1 and phase 2 enzymes, cuticular proteins, and receptor sites. Examination of down-regulated transcripts, including receptor sites (none of which were observed to be statistically down-regulated) was conducted for each experimental group, however our efforts were primarily focused on the classification of over-expressed, enzymatic detoxification transcripts which could result in the production of more proteins, and could theoretically increase the detoxification of the pesticide insult. A search within each set of differentially-expressed transcripts was conducted. While we observed multiple detoxification mechanisms as a response to imidacloprid and chlorothalonil exposure, we did not observe any known detoxification mechanisms as a result of exposure to boscalid. This observation is interesting because it has been previously noted that boscalid can have a driving selection on this insect taxa^28^. More detailed examination of the detoxification mechanisms in common between the imidacloprid and chlorothalonil treatment groups revealed two enzymatic mechanisms of detoxification: a cytochrome p450 (**6k1 isoform X1)** and a UDP-glucuronsyltransferase (**2b7-like**). Several cytochrome p450 enzymes have previously been implicated in imidacloprid resistance in *L. decemlineata*, including in a study performed by Kaplanogula et al. (2017) where the over-expression of cytochrome p450 4Q3 and UDP-glucuronsyltransferase 2 was linked to imidacloprid resistance^18^. The over-expression of the two similar enzymatic detoxification mechanisms described within our study suggests exposure to the fungicide chlorothalonil could induce a similar genetic response to the insecticide imidacloprid, which could eventually lead to increased levels of insensitivity to other pesticide insults, and the potential for cross-resistance in a field setting. An additional and important distinction centers on the number of dissimilar mechanisms between both imidacloprid and chlorothalonil, suggesting that while we did observe similarities, the detoxification and genetic responses to these pesticides are also quite unique.

When a correlation matrix was generated from the transcriptomic and RNA sequencing data, a grouping of three individuals (Imidacloprid_2, Imidacloprid_3 and Chlorothalonil 1) was clearly observed, and the expression patterns of these three insects were statistically similar to each other. Differentially-expressed transcripts between these individuals and the remainder of the test insects was further evaluated. This examination revealed highly over-expressed detoxification mechanisms including acetylcholine receptor subunit (alpha-like), cytochrome p450’s, cuticular proteins, multi-drug resistance proteins and an oxidoreductase subunit. Further, to determine whether the transcriptomic expression of these three individuals was based on some biological process and not the response of pesticide exposure, we examined the transcript expression of these individuals for a multitude of several biological processes including spermatic transcripts (could suggest a sex-linked factor) and apoptotic genes. Further, examination of the over-expressed transcripts’ GO terms was also conducted. Taken together, these supplemental examinations revealed similarities in over-expression of sperm related transcripts including transcripts encoding for spermatogenesis, suggesting that these individuals were probably all male at the same developmental stage following emergence from pupation. No apoptotic genes were found and over expression of iron production and oxidoreducatase activity in the GO terms was noted. Overall, the genetic analysis suggested that all three insects (e.g. Imidacloprid_2, imidacloprid_3 and Chlorothalonil_1) responded similarly to their pesticide exposure.

To confirm the results from edgeR and the differentially-expressed transcripts, expression patterns were confirmed using qPCR. The expression determined by qPCR was very similar to what edgeR produced, validating the differentially-expressed data. To further confirm the importance of the two similar genetic detoxification mechanisms (cytochrome p450 6k1 isoform X1 and a UDP-glucuronsyltransferase 2b7-like) in imidacloprid and chlorothalonil detoxification, we re-evaluated our pesticide feeding assays independently using two additional replicate assays and measured the expression of our two transcripts of interests. From these two additional replicate assays, we observed high variation in transcript expression between replicates for both cytochrome p450 6k1 isoform X1 and the UDP-glucuronsyltransferase 2b7-like. Multiple individuals over-expressed both genetic mechanisms in both exposure groups, while other individuals also showed no expression differences. The average of the biological replicates is presented in Table 6 and their corresponding CT values with corresponding STD is presented in **Supplemental Table S3**. Within the two additional chlorothalonil replicates, we observed over-expression of both cytochrome p450 6k1 isoform X1 and a UDP-glucuronsyltransferase 2b7-like transcripts, whereas in the two additional imidacloprid replicates we observed only slight over-expression of the UDP-glucuronsyltransferase in one group and no over-expression of any other mechanisms in any other group. These findings further suggest that the activation of putative resistance mechanisms is highly variable between individuals, but are partially regulated by pesticide exposure and should be further examined in field relevant populations for their role in insecticide resistance.

The aim of our transcriptomic study was to classify the genetic detoxification mechanisms activated from exposure of the insecticide imidacloprid and the fungicides chlorothalonil and boscalid, and further to determine if there was any overlap in the genetic response within these individuals. In our exploration we found several over-expressed transcripts as a response to these chemicals. Further, we were able to classify two enzymatic detoxification mechanisms activated in common by both imidacloprid and chlorothalonil. Interestingly, individual beetle responses varied greatly to their chemical stressors, suggesting that these genetic responses, while similar, could also be dependent upon individual genetics. Examination of the genetic response imposed by commonly occurring pesticides on both target and off-target individuals can lead to a better understanding of how insecticide resistance develops. While this study was conducted with a naïve lab colony, future studies from field relevant individuals would help confirm the significance of our results and the effects of off-target pesticide treatments.

## Acknowledgements

The authors would like to acknowledge support from the Wisconsin Potato and Vegetable Growers Association and the associated agricultural community. We would also like to acknowledge support from the University of Wisconsin-Madison Bioinformatics Research Center. This research was partially supported by a combination of funding from: (1) USDA-NIFA-ARFI-ELI Postdoctoral Fellowship Grant 12105364 and (2) support from the Wisconsin Potato and Vegetable Growers Association.

